# Pluripotency and the origin of animal multicellularity

**DOI:** 10.1101/564518

**Authors:** Shunsuke Sogabe, William L. Hatleberg, Kevin M. Kocot, Tahsha E. Say, Daniel Stoupin, Kathrein E. Roper, Selene L. Fernandez-Valverde, Sandie M. Degnan, Bernard M. Degnan

**Author notes:** These authors contributed equally to this work (Shunsuke Sogabe, William L. Hatleberg). The Scottish Oceans Institute, Gatty Marine Laboratory, School of Biology, University of St Andrews, East Sands, St Andrews, Fife KY16 8LB, UK (S.S.); Department of Biological Sciences, Carnegie Mellon University, 4400 Fifth Avenue, Pittsburgh, PA 15213 USA (W.L.H.); BioQuest Studios, PO Box 603, Port Douglas QLD 4877, Australia (D.S.); Centre for Clinical Research, Faculty of Medicine, University of Queensland, Herston QLD 4029, Australia (K.R.); CONACYT, Unidad de Genómica Avanzada, Laboratorio Nacional de Genómica para la Biodiversidad, Centro de Investigación y de Estudios Avanzados del IPN, Irapuato, Guanajuato, Mexico (S.L.F.-V.).

## Abstract

The most widely held, but rarely tested, hypothesis for the origin of animals is that they evolved from a unicellular ancestor with an apical cilium surrounded by a microvillar collar that structurally resembled present-day sponge choanocytes and choanoflagellates^1–4^. Here we test this traditional view of the origin of the animal kingdom by comparing the transcriptomes, fates and behaviours of the three primary sponge cell types – choanocytes, pluripotent mesenchymal archeocytes and epithelial pinacocytes – with choanoflagellates and other unicellular holozoans. Unexpectedly, we find the transcriptome of sponge choanocytes is the least similar to the transcriptomes of choanoflagellates and is significantly enriched in genes unique to either animals or to sponges alone. In contrast, pluripotent archeocytes upregulate genes controlling cell proliferation and gene expression, as in other metazoan stem cells and in the proliferating stages of two closely-related unicellular holozoans, including a colonial choanoflagellate. In the context of the body plan of the sponge, *Amphimedon queenslandica*, we show that choanocytes appear late in development and are the result of a transdifferentiation event. They exist in a metastable state and readily transdifferentiate into archeocytes, which can differentiate into a range of other cell types. These sponge cell type conversions are similar to the temporal cell state changes that occur in many unicellular holozoans^5^. Together, these analyses offer no support for the homology of sponge choanocytes and choanoflagellates, nor for the view that the first multicellular animals were simple balls of cells with limited capacity to differentiate. Instead, our results are consistent with the first animal cell being able to transition between multiple states in a manner similar to modern transdifferentiating and stem cells.

## Main

The last common ancestor of all living animals appears to have possessed epithelial and mesenchymal cell types that could transdifferentiate over an ontogenetic life cycle (Fig.1a)^1,4^. This capacity to develop and differentiate required a regulatory capacity to control spatial and temporal gene expression, and included a diversified set of signalling pathways, transcription factors, enhancers, promoters and non-coding RNAs (Fig. 1a)^5–9^. Recent analyses of the genomes and life cycles of unicellular holozoan relatives of animals have revealed that the regulatory repertoire present in multicellular animals largely evolved first in a unicellular ancestor (Fig. 1a)^2,5,6^. These insights contrast with a widely-held view that all animals evolved from a stem organism that was a simple ball of ciliated cells^1,3,4^. Implicit in this traditional perspective is that (i) regulatory systems necessary for cell differentiation evolved after the divergence of metazoan and choanoflagellates lineages, and (ii) morphological features shared between choanoflagellate and choanocytes are homologous and were present in the original animal cell. While the former is not supported by recent data - unicellular holozoans can change cell states by environmentally-induced temporal shifts in gene expression (Fig. 1a)^5,6,10–12^ – the latter is contingent upon the still controversial aspect of whether extant choanocytes and choanoflagellates accurately reflect the ancestral animal cell type.

To test this, we first compared cell type-specific transcriptomes^13^ from the sponge *Amphimedon queenslandica* with each other, and with transcriptomes expressed during the life cycles of closely-related unicellular holozoans, the choanoflagellate *Salpingoeca rosetta*, the filasterean *Capsaspora owczarzaki* and the ichthyosporean *Creolimax fragrantissima* (Fig. 1a)^10–12^. We chose three sponge somatic cell types hypothesised to be homologous to cells present in the last common ancestor of contemporary metazoans, choanozoans or holozoans: (i) choanocytes, which are internal epithelial feeding cells that capture food by pumping water through the sponge; (ii) epithelial cells called pinacocytes, which line internal canals and the outside of the sponge; and (iii) mesenchymal pluripotent stem cells called archeocytes, which inhabit the middle collagenous layer and have a range of other functions (Fig. 1 and Supplementary Video 1)^2,14–16^. These three cell types were manually picked and frozen within 15 minutes of *A. queenslandica* being dissociated (see Methods and Supplementary Video 2). Their transcriptomes were sequenced using CEL-Seq2^17^ and mapped to the Aqu2.1 annotated genome^18^. This approach allowed visual verification of the three cell types, minimised the time for transcriptional changes to occur after cell dissociation, and allowed for deep sequencing of cell type transcriptomes (Extended Data Table 1, and Supplementary Files S1 and S2).

**Figure 1.**
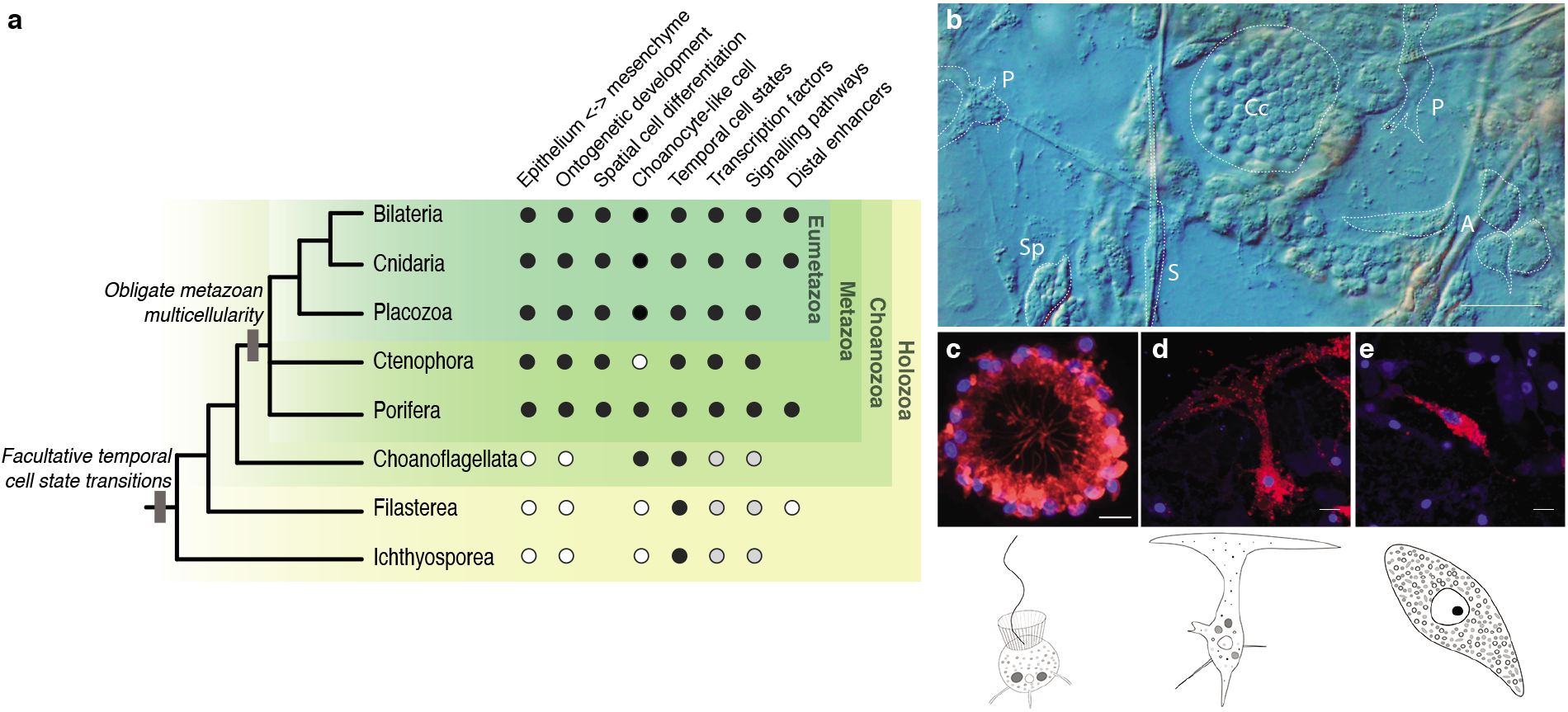
Holozoan relationships and sponge cell types. **a,** Cellular and regulatory traits of metazoans and closely related unicellular holozoans. Black dots, trait present; white dots, trait absent; grey dots, trait present but to a lesser extent. Two major evolutionary events are mapped onto the holozoan phylogenetic tree: (i) environmentally-induced, facultative changes in cell state ancestral to holozoans; and (ii) obligate metazoan multicellularity. **b,** Whole mount internal view of a juvenile *Amphimedon queenslandica*. Cell types are outlined. A, archeocyte (cluster of four outlined); Cc, choanocyte chamber; S, sclerocyte; Sp, spherulous cell; P, pinacocyte. **c,** Choanocyte chamber labelled with DiI with an illustration of a single choanocyte below. **d,** Pinacocyte labelled with DiI with illustration below. **e,** Archeocyte labelled with DiI with illustration below. Scale bars: b, 10 μm; c-e, 5 μm.

Principle component analysis (PCA) and sparse partial least squares discriminant analysis (sPLS-DA)^19^ reveal that the transcriptomes of the three *A. queenslandica* cell types are unique, with choanocytes being the most distinct (Fig. 2 a and Extended Data Fig. 1). Of 44,719 protein-coding genes, 11,013 genes were identified as significantly differentially expressed in at least one cell type from pairwise comparisons between the three cell types using DESeq2^20^ (Fig. 2b and Supplementary File S3). Significant differences between cell types were independently corroborated by sPLS-DA, which highlighted a subset of 110 genes that explain 15% of the variance in the dataset and clearly discriminate the choanocytes from the other two cell types (Extended Data Fig. 1). This subset includes numerous putative immunity genes that typically encode multiple domains in unique configurations, including scavenger receptor cysteine-rich, tetratricopeptide repeat and epidermal growth factor domains (Supplementary File S4).

**Figure 2.**
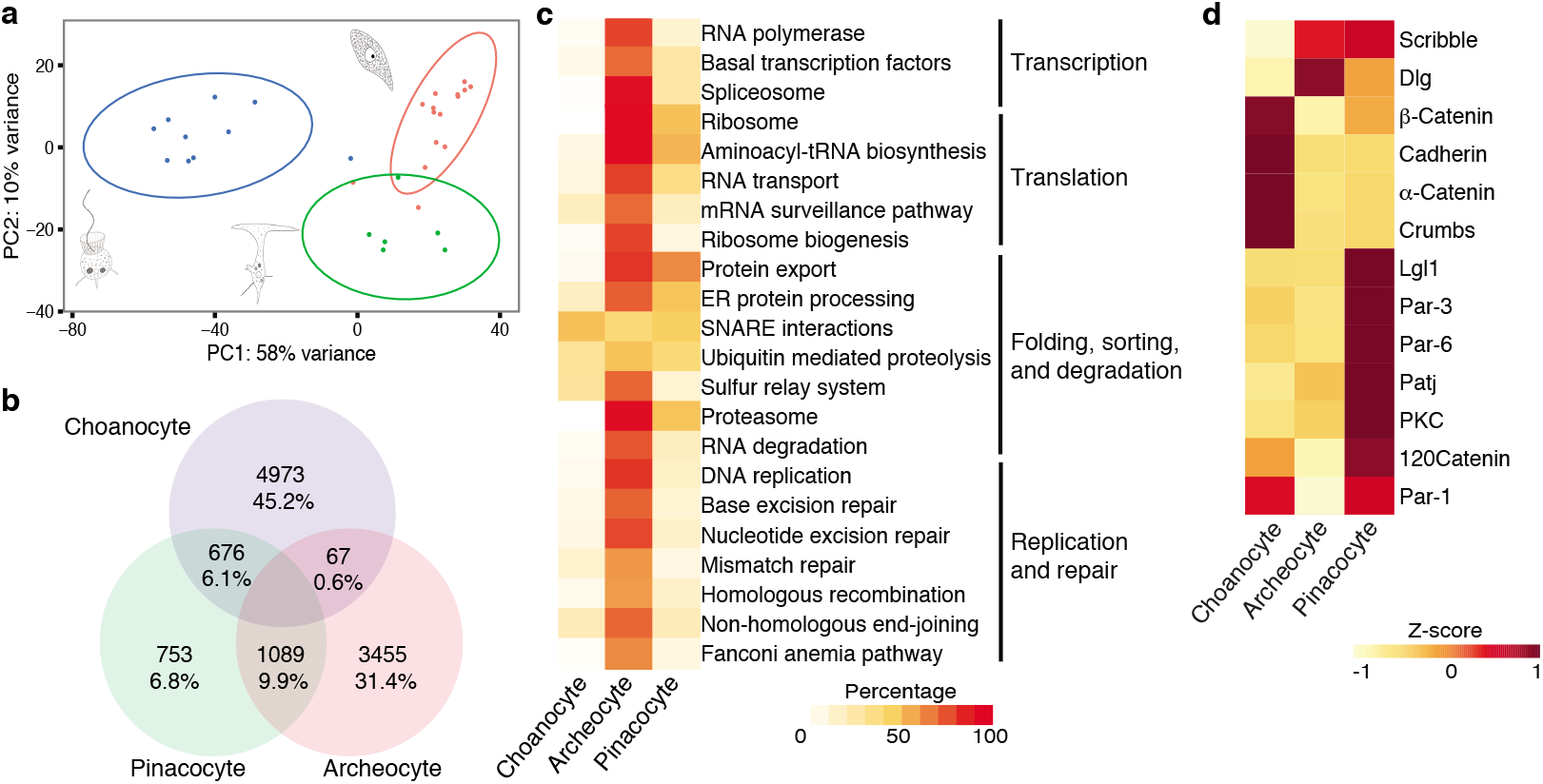
Comparison of choanocyte, archeocyte and pinacocyte transcriptomes. **a,** PCA plot of CEL-Seq2 transcriptomes with 95% confidence level ellipse plots. Blue, choanocytes; red, archeocytes; green, pinacocytes. **b,** Venn diagram summary of the number of significantly up-regulated genes based on pairwise comparisons between each of the three cell types using DESeq2 with a false discovery rate (FDR) < 0.05. The percentages are of the total genes differentially up-regulated in all cell types. **c,** Percentage of KEGG Genetic Information Processing genes present in each cell type, corresponding to the number of components making up each KEGG category identified. **d,** Heat map of the expression of *Amphimedon* epithelial cell polarity, junction and basal lamina genes in each cell type.

From the DESeq2 analysis, we find that archeocytes significantly upregulate genes involved in the control of cell proliferation, transcription and translation, consistent with their function as pluripotent stem cells (Fig. 2c and Supplementary File S5). In contrast, choanocyte and pinacocyte transcriptomes are enriched in suites of genes involved in cell adhesion, signalling and polarity, consistent with their role as epithelial cells (Fig. 2d; Extended Data Figure 2 and Supplementary File S5).

We identified the evolutionary age of all protein coding genes in the *Amphimedon* genome as well as the genes significantly and uniquely up-regulated in each cell-type specific transcriptome using phylostratigraphy, which is based on sequence similarity with genes in other organisms with a defined phylogenetic distance^21^. Specifically, we classified *Amphimedon* genes as having evolved (i) before or (ii) after divergence of metazoan and choanoflagellate lineages (these are called pre-metazoan and metazoan genes, respectively), or (iii) after divergence of the sponge lineage from all other animals (sponge-specific genes). In total, the *A. queenslandica* genome is comprised of 28% pre-metazoan, 26% metazoan and 46% sponge-specific protein-coding genes (Fig. 3a and Supplementary File S6). We find that 43% of genes significantly up-regulated in choanocytes have homologues detectable only in sponges, which is similar to the entire genome (Fig. 3b). In contrast, 62% of genes significantly up-regulated in the pluripotent archeocytes belong to the evolutionarily oldest pre-metazoan category, which is significantly higher than 28% for the entire genome (Fig. 3c). As with archeocytes, pinacocytes express significantly more pre-metazoan and fewer sponge-specific genes than would be expected from the whole genome profile (Fig. 3d). Results supporting this analysis are obtained when we (i) undertake the same phylostratigraphic analysis of all genes expressed in these cell types, taking also into account relative transcript abundances (Extended Data Fig. 3 and Supplementary File S7), or (ii) classify gene age using an alternative method to identify of orthogroups (homology cluster containing both orthologues and paralogues)^22^ among unicellular holozoan, yeast and *Arabidopsis* coding sequences (Extended Data Fig. 4).

**Figure 3.**
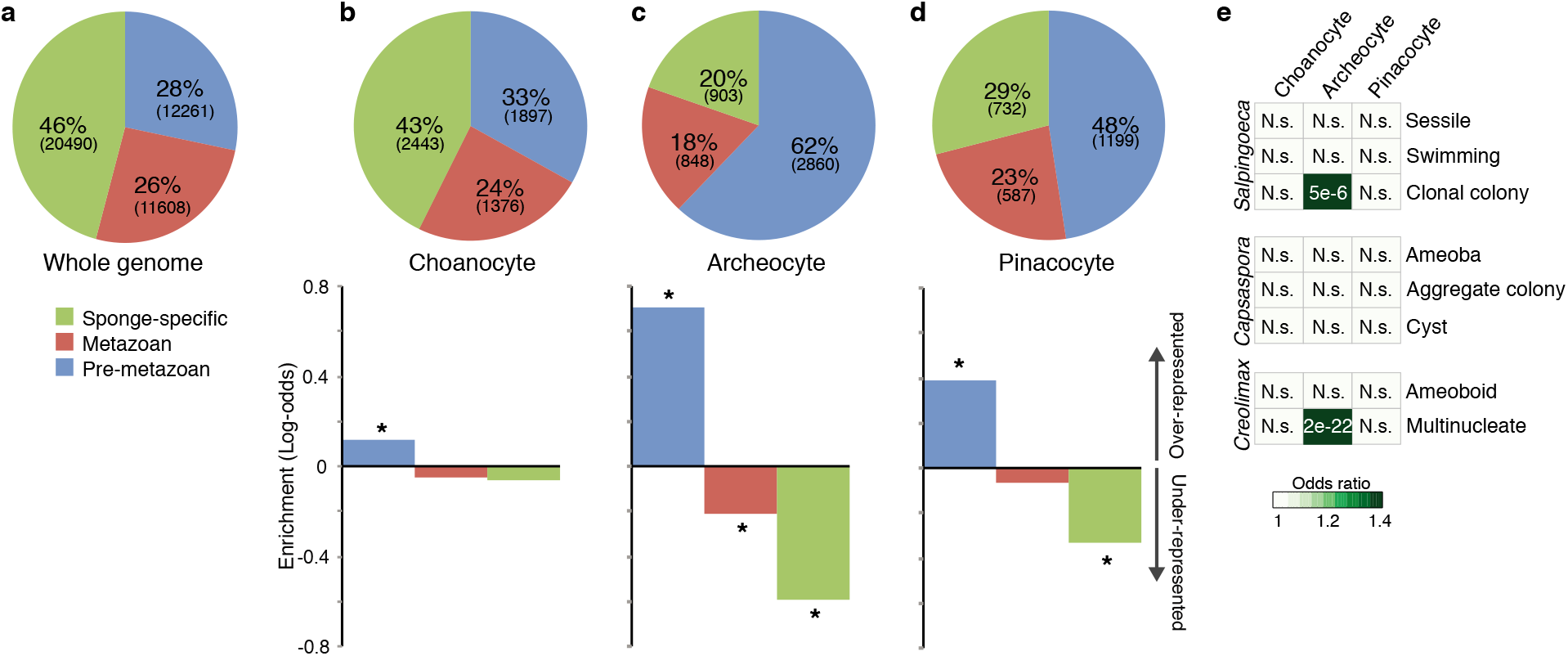
Analysis of gene age of choanocyte, archeocyte and pinacocyte transcriptomes. **a,** Phylostratigraphic estimate of the evolutionary age of coding genes in the *A. queenslandica* genome. **b-d,** Estimate of gene age of differentially-expressed genes in choanocytes (b), archeocytes (c) and pinacocytes (d) and the enrichment of phylostrata relative to the whole genome (bottom). Asterisks indicate significant difference (p-value <0.001) from the whole genome. **e,** A heat map comparing orthologous genes uniquely up-regulated in *A. queenslandica* cell types, and *Salpingoeca rosetta, Capsaspora owczarzaki* and *Creolimax fragrantissima* life cycle stages. Colour indicates the significance of overlap based on the odds ratio. Values indicate adjusted p-values and show significant resemblance only between the archeocyte and the *S. rosetta* colonial stage and the *C. fragrantissima* multinucleate stage. N.s., not significant.

Comparison of *A. queenslandica* cell-type transcriptomes with stage-specific transcriptomes from the choanoflagellate *S. rosetta*^10^, the filasterean *C. owczarzaki*^11^ and the ichthyosporean *C. fragrantissima*^12^ reveals that archeocytes have a significantly similar transcriptome to the colonial stage of the choanoflagellate and the multinucleate stage of the ichthyosporean (Fig. 3e). Consistent with this result, the significantly up-regulated genes in the colonial or multinucleate stages of all three unicellular holozoans share the highest proportion of orthogroups with genes significantly up-regulated in archeocytes (Extended Data Fig. 5). In contrast, choanocyte and pinacocyte transcriptomes have no significant similarity to any known unicellular holozoan transcriptome, and share a lower proportion of orthogroups with unicellular holozoans compared to archeocytes (Fig. 3e and Extended Data Fig. 5a).

When we compare the 94 differentially up-regulated transcription factor genes in *A. queenslandica* choanocytes, pinacocytes and archeocytes, we find no marked difference in their phylostratigraphic age, suggesting that the gene regulatory networks operational in these cells are of an overall similar evolutionary age (Extended Data Fig. 6 and Supplementary File S8). We detected 20, 25 and 21 orthologues of the 43 evolutionarily-oldest (i.e. pre-metazoan) transcription factor genes expressed in the *Amphimedon* cells in the genomes of *Salpingoeca, Capsaspora* and *Creolimax* respectively, with 9 of these being present in all species (Supplementary File S8). Comparison of the expression profiles of the transcription factor genes shared among these unicellular holozoans and *Amphimedon* revealed no evidence of a conserved, coexpressed gene regulatory network (Extended Data Fig. 7 and Supplementary File S8). However, the proto-oncogene *Myc* and its heterodimeric partner *Max* are up-regulated in *A. queenslandica* archeocytes (Extended Data Fig. 6), as observed in other metazoan self-renewing pluripotent stem cells^23^. Myc and Max are present also in choanoflagellates, filastereans and ichthyosporeans, where they heterodimerise and bind to E-boxes just as they do in animals^10–12,24^. Myc is expressed in the proliferative stage of *Capsaspora*, where it regulates genes associated with ribosome biogenesis and translation^6^. Sponge archeocytes also have enriched expression of genes involved in translation, transcription and DNA replication (Fig. 2c). This suggests that Myc’s role in regulating proliferation and differentiation predates its role in bilaterian stem cells and cancer^23,25^, and was likely a cardinal feature of the first metazoan cell.

Given that we found no transcriptional support for homology of *A. queenslandica* choanocytes and choanoflagellates, but did find evidence for pluripotent archeocytes expressing a largely premetazoan transcriptome, we sought to investigate the relationships of these cell types in the context of development and the body plan. In *Amphimedon* and most other sponges, archeocytes form during embryogenesis to populate the inner cell mass of the larva and are the most prevalent cell type during early metamorphosis^15^. As metamorphosis progresses, these archeocytes differentiate into other cell types that populate the juvenile body plan, including choanocytes and pinacocytes, the former of which can transdifferentiate into other cell types^16,26^. To further understand the stability of choanocytes and the dynamics of transdifferentiation, we selectively labelled choanocytes in 3 day old juvenile *A. queenslandica* with CM-DiI (Fig. 4a) and followed their fate over 24 hours (Fig. 4b). Within 4 hours of labelling, many choanocytes dedifferentiated into archeocytes (Fig. 4c, d, Supplementary Video 3); this did not require prior cell division (Extended Data Fig. 8). By as little as two hours later, some of these CM-DiI labelled archeocytes had differentiated into pinacocytes (Fig. 4e); within 12 hours, we detected multiple labelled cell types (Fig. 4e, f). Together, these results suggest that archeocytes are essential in the development and maintenance of the *A. queenslandica* body plan, as appears to be the case in other sponges^15^. Unlike archeocytes, choanocytes appear late in development and exist in a metastable state, sometimes lasting only a few hours before dedifferentiating back into archeocytes (Fig. 4g, Extended Data Fig. 8).

**Figure 4.**
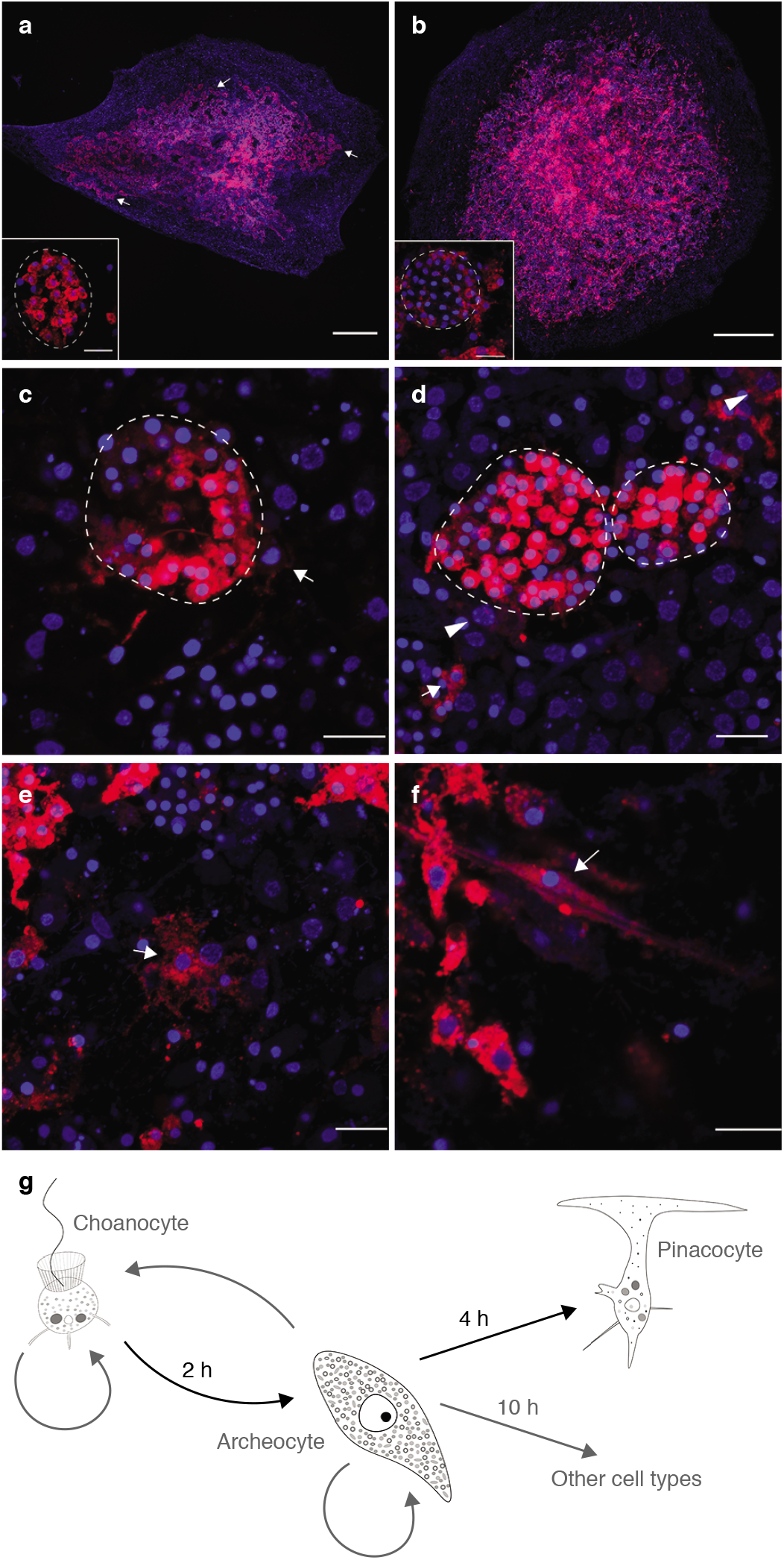
Transdifferentiation of choanocytes in *Amphimedon queenslandica*. **a, b,** Whole mount views of 4 day old juveniles labelled with CM-DiI. a, 30 min after CM-DiI labelling. Labelling is almost exclusively in choanocytes in chambers; insert, a single labelled choanocyte chamber. **b,** 24 hours after labelling. CM-DiI labelling spread throughout the juvenile with limited staining still present in choanocyte chambers; insert, a choanocyte chamber comprising largely of unlabelled cells. **c, d,** 2 hours (c) and 4 hours (d) after labelling. Labelled cells (arrow) migrating outside of choanocyte chambers (dotted lines), some of which have a large nucleus and a clearly visible nucleolus (arrowheads) characteristic of archeocytes. **e,** 6 hours after initial DiI labelling of choanocytes, labelled pinacocytes (arrow) with thin pseudopodia are detected. **f,** 12 hours after initial labelling, CM-DiI labelled skeletal sclerocytes (arrow) and other cell types are present. **g,** Summary diagram of cell type transition in the *A. queenslandica* juvenile. Scale bars: a, b, 200 μm; c-f, 10 μm.

In conclusion, our analysis of sponge and unicellular holozoan cell transcriptomes, development and behaviour provides no support for the long-standing and widely-held hypothesis that multicellular animals evolved from an ancestor that was an undifferentiated ball of cells resembling extant choanocytes and choanoflagellates^1–4^. This conclusion is corroborated by recent studies that question the homology of choanocytes and choanoflagellates based on cell structure^27,28^. As an alternative, we posit that the ancestral metazoan cell type, regardless of its external character, had the capacity to exist in, and transition between, multiple cell states in a manner similar to modern transdifferentiating and stem cells. Previous analyses of holozoan genomes support this postulate, with some of the genomic foundations of pluripotency being established deep in a unicellular past^6,24^. Genomic innovations unique to metazoans, including the origin and expansion of key signalling pathway and transcription factor families, and regulatory DNA and RNA classes^7,9,29^, may have conferred the ability of this ancestral pluripotent cell to evolve a regulatory system where it could co-exist in multiple states of differentiation, giving rise to the first multicellular animal.

**Supplementary Information** is linked to the online version of the paper at www.nature.com/nature. (This submission includes eight Supplementary Information data files as well as additional material that is available on Dryad.)

## Acknowledgements

This study was supported by funds from the Australian Research Council (B.M.D. and S.M.D.). We thank Iñaki Ruiz Trillo for primary expression data for *Capsaspora* and *Creolimax*.

## Author Contributions

B.M.D and S.M.D conceived and designed the project. S.S., D.S. and K.R. identified and isolated the cells, and prepared the libraries. W.H., S.S and K.M.K. undertook gene expression and annotation, and phylostratigraphic analyses with help from T.S., S.M.D, S. F.-V and B.M.D. S.S. undertook cell lineage analyses. B.M.D, S.M.D and S.S. wrote the manuscript with comments and contributions from all authors.

## Author Information

### Data deposition statement

*Amphimedon queenslandica* genome sequence can be accessed at (http://metazoa.ensembl.org/Amphimedon_queenslandica/Info/Index).

All cell-type transcriptome data are available in the NCBI SRA database under the BioProject PRJNA412708. Additional supplementary data is `available upon request`

## Methods

### Cell isolation

Adult *Amphimedon queenslandica* were collected from Heron Island Reef, Great Barrier Reef and transferred to a closed aquarium facility where they were housed for no more than three days before being cut into approximately 1 cm^3^ cubes. These cubes were mechanically dissociated by squeezing through a 20 μm mesh. The resultant cell suspension was diluted with 0.22 μm-filtered seawater (FSW) and the target cell types were identified microscopically based on morphology. Archeocytes are much larger than the other cells and possess a highly visible nucleolus. Choanocytes remain in intact choanocyte chambers after dissociation. Pinacocytes, unlike the other cell types, are translucent and maintain protruding cytoplasmic processes after dissociation. This approach avoided misidentification of dissociated cell types, but could not determine whether these cells are in the process of dividing or differentiating. Individual cells or choanocyte chambers were collected under an inverted microscope (Nikon Eclipse Ti microscope) using a micropipette mounted on micromanipulator (MN-4, Narishige) connected to CellTram Oil (Eppendorf) (Supplementary video 2), flash frozen and stored at −80°C. All cells were frozen within 15 min of dissociation. Samples used in CEL-Seq2 were comprised of pools of either five to six archeocytes or pinacocytes, or a single choanocyte chamber (~40-60 cells) (Extended Data Table 1). Based on differences in cell size, we estimated that these pools have similar amounts of total RNA. Three pinacocyte, and five archeocyte and choanocyte samples were collected from each of three sponges (Supplementary File S2).

### CEL-Seq2 sample preparation, sequencing and analysis

Samples were prepared according to the CEL-Seq2 protocol^17^ and sequenced on two lanes of Illumina HiSeq2500 on rapid mode using HiSeq Rapid SBS v2 reagents (Illumina); CEL-Seq2 libraries were randomised in relation to cell type and source adult sponge in these two lanes. CEL-Seq2 reads were processed using a publicly available pipeline (https://github.com/yanailab/CEL-Seq-pipeline;see additional supplementary data on Dryad:/CEL-Seq pipeline/). Read counts were obtained from demultiplexed reads mapped to *A. queenslandica* Aqu2.1 gene models^18^. Samples with read counts less than 10^6^ were removed and not included in subsequent analyses (Supplementary File S2). For the samples included in the final analysis, approximately 60% of the reads successfully mapped to the genome (Extended Data Table 1), as per other studies using CEL-Seq^30^.

### Analysis of differentially expressed genes

The mapped read counts were analysed for differential gene expression using the bioconductor package DESeq2^20,31^ (<u>see additional supplementary data on Dryad: /DESeq2/</u>). Genes that had read counts with a row sum of zero were removed. Principle component analyses (PCA) were performed on blind variance stabilising transformed (vst) counts obtained using DESeq2 and were visualised using the ggplot2 package^32^. Pairwise comparisons were conducted between each of the three cell types to generate a differentially expressed gene (DEG) list for each cell type using a false discovery rate (FDR) < 0.05. Venn diagrams were generated using VENNY (http://bioinfogp.cnb.csic.es/tools/venny) to visualise and compare the list of DEGs between each cell type. Heat maps were generated using the R-packages pheatmap^33^ and RColorBrewer^34^ to visualise the expression patterns between the cell types using the vst transformed counts, which were scaled into z score values ranging from -1 (low expression) to 1 (high expression).

All protein coding genes were annotated using blastp (e-value cutoff = 1e-3) and InterProScan (default settings), which were merged in Blast2GO^35,36^. KEGG annotations were obtained using the online tool BlastKOALA^37^ (<u>see additional supplementary data on Dryad: /KEGG annotation)</u>. Pathway analyses were performed using the annotations on the KEGG Mapper - Reconstruct Pathway tool^38^. Complete DEG lists with BLAST2GO, InterPro, Pfam, and phylostrata ID can be found in Supplementary File S3, as well as KEGG pathway enrichments in Supplementary File S5.

To identify the genes that best explain differences among cell type transcriptomes, we adopted the multivariate sparse Partial Least Squares Discriminant Analysis (sPLS-DA)^19^, implemented in the mixOmics package^39^ in R v3.3.1 (<u>see additional supplementary data on Dryad: /sPLS-DA/README.txt</u>). This is a supervised analysis that uses the sample information (cell type) to identify the most predictive genes for classifying the samples according to cell type. The optimised numbers of genes per component were obtained by training and correctly evaluating the performance of the predictive model using 5-fold cross-validation, repeated 100 times. A sample plot was used to visualise the similarities between samples for the final sPLS-DA model with 95% confidence ellipses using the plotIndiv function in R. A heat map was used to visualise relative expression levels of the selected gene models for the two components, using vst counts and the package pheatmap^33^ in R. Venn diagrams were generated using VENNY to visualise and compare the DEGs generated by DESeq2 and sPLS-DA.

### Phylostratigraphy

To estimate the evolutionary age of genes up-regulated in each cell type, phylostratigraphy analyses^21^ were performed using blastp and an e-value cutoff of 0.001 on a custom database containing 1,757 genomes and transcriptomes^40^ that was modified to account for *A. queenslandica*’s phylogenetic position (i.e. all eumetazoan and bilaterian taxa were moved into the metazoan phylostratum, and three phylostrata - poriferan, demosponge and haplosclerid - were added to increase the representation of poriferan transcriptomes; Supplementary File S6, <u>see additional supplementary data on Dryad: /Phylostratigraphy annotations/</u>). Every gene model in *A. queenslandica* was blasted against each sequence in the database, and its age of gene origin was inferred based on the oldest blast hit relative to a predetermined phylogenetic tree (<u>see additional supplementary data on Dryad: /Phylostratigraphy annotations/</u>).

Phylostrata enrichments were performed using the Fisher’s exact test^41^ in the BioConductor package, GeneOverlap^42^ in R, to identify significant differences in gene age of the cell type DEG lists relative to the genome (<u>see additional supplementary data on Dryad</u>: /Fig.3b-d and /ED_Fig3_files). Enrichment (log odds ratio value above 0) and under-representation (log odds ratio value below 0) of each phylostrata found in the cell type DEG lists relative to the genome, were visualised using the R-packages pheatmap^33^ and RColorBrewer^34^.

### Orthology analyses

Orthology analyses were performed using FastOrtho^43^ from a custom ‘all-vs-all’ blastp database of coding sequences from the genomes of *Saccharomyces cerevisiae^44^, Arabidopsis thaliana^45^, Creolimax fragrantissima^12^, Sphaeroforma arctica^46^, Capsaspora owczarzaki^47^, Monosiga brevicollis^48^*, and *Salpingoeca rosetta^10^*, using the following configuration settings: pv_cutoff = 1e-5; pi_cutoff = 0.0; pmatch_cutoff = 0.0; maximum_weight = 316.0; inflation = 1.5; blast_e = 1e-5 (<u>see additional supplementary data on Dryad</u>: /FastOrtho/). FastOrtho classifies all of the genes present in each genome into orthology groups (orthogroups, OGs), which contain all orthologous and paralogous genes from each species. Genes that do not have any orthologues in other species or paralogues within the same genome were not included in any orthogroups. To compare the gene lists between species in all downstream analyses, species-specific gene names were changed to the common orthogroup identifier.

Orthology analyses between *A. queenslandica* and *S. rosetta, C. fragrantissima*, and *C. owczarzaki* cell types were performed using the cell type-specific DEG lists obtained from previous studies on *S. rosetta*^10^, *C. fragrantissima*^12^, and *C. owczarzaki*^11^. The BioConductor package, GeneOverlap^42^, was used to identify (1) the number overlapping OGs between species and cell type, and (2) the statistical significance of that overlap based on list size and total number of OGs (<u>see additional supplementary data on Dryad</u>: /Fig.3e). This function provided the odds ratio between the OG lists, where the null hypothesis was no significant overlap (odds ratio value of 1 or smaller) and the alternative being a significant overlap detected between the lists (odds ratio value over 1), as well as a p-value calculated for odds ratio values over 1.

To supplement phylostratigraphy analyses of *Amphimedon* cell-type specific gene lists (Fig. 3 and Extended Data Fig. 3), the BioConductor package, GeneOverlap^42^ was used to identify the number and percentage of orthogroups that are also present in the genomes of *Arabidopsis thaliana, Saccharomyces cerevisiae, Creolimax fragrantissima, Sphaeroforma arctica, Capsaspora owczarzaki, Monosiga brevicollis, and Salpingoeca rosetta* (Extended Data Fig. 4 and Extended data Fig. 5; <u>see additional supplementary data on Dryad</u>: /ED_Fig4 and ED_Fig5)

### Classification of gene expression levels into quartiles

In addition to differential gene expression analyses for *Amphimedon* transcriptomes, the relative gene expression levels for all cell types were assigned to one of four expression quartiles based on the number of reads that mapped to a given Aqu2.1 gene model (Extended data Fig. 3). All zero read counts were discarded and the mean expression value of the non-transformed normalised count values of all samples (from all cell types) was used to calculate the quartile values. These values (Q1: 2.30, Q2: 6.06, Q3: 15.83) were used to classify the expression of all of genes in each cell type into four groups based on transcript abundance, ranging from lowest (Q1) to highest (Q4).

Phylostrata enrichments for the different quartile value thresholds were performed as described above for the cell type DEG lists; heat maps were generated using pheatmaps^33^ in R (<u>see additional supplementary data on Dryad</u>: /ED_Fig3_files). All downstream analyses used the median value (Q2: 6.06) as a cut-off value to obtain a list of expressed genes. Orthology analyses using FastOrtho were performed as described above, and the percentage of genes with shared orthologous group (OG) in each gene list was calculated (<u>see additional supplementary data on Dryad</u>: /ED_Fig4_files and ED_Fig5_files). In these analyses, exclusive lists refer to all of the regions in the Venn diagram being treated as a separate list (e.g. archeocyte only, common between archeocyte and choanocyte, common between archeocyte and pinacocyte, etc.), while non-exclusive lists collapse all of the lists containing a given cell type into one list (e.g. archeocyte non-exclusive DEG list includes, archeocyte DEGs + (archeocyte + pinacocyte DEGs) + (archeocyte + choanocyte DEGs).

### Identification and analysis of expressed *A. queenslandica* transcription factors

A list of *A. queenslandica* transcription factors expressed in the three cell types was obtained using a number of independent methods. First, a non-conservative list of putative *A. queenslandica* transcription factors was obtained using the DNA-binding domain database (DBD: Transcription factor prediction database) and the Pfam IDs of sequence specific DNA-binding domain (DBD) families, which corresponds to known transcription factor families (www.transcriptionfactor.org^49^). Second, we collated a list of annotated *A. queenslandica* transcription factors in the literature^7,16,47,50–66^ (Supplementary File S8). Third, we compared these lists to an unpublished in-house database for *A. queenslandica* (Degnan *et al*. unpublished) and putative transcription factors identified by OrthoMCL. The final list of 173 expressed transcription factor genes used in this study were present in at least two of the three lists (Supplementary File S8).

The evolutionary age of each of the expressed transcription factors was first assigned based on the DBD contained in the gene model and then manually curated based primarily on literature (Supplementary File S8). From this, each TF was assigned as either originating in sponges after diverging form other animals (sponge-specific), in metazoans after they diverged from choanoflagellates (metazoan) or before metazoans diverged from choanoflagellates (premetazoan).

### Analysis of juvenile cell fate and proliferation

Larvae were collected as previously described^67^, left in FSW overnight and then placed in sterile 6-well plates with 10 ml of FSW for 1 hour in the dark with live coralline algae *Amphiroa fragilissima*. Postlarvae settled on *A. fragilissima* were removed using fine forceps (Dumont #5) and resettled on to round coverslips placed in a well with 2 ml FSW in a sterile 24-well plastic plate, with 3 postlarvae placed on each coverslip. Metamorphosis from resettled postlarvae to a functional juvenile takes approximately 72 hours^16,68^. For all samples, FSW was changed daily until fixation.

The lipophilic cell tracker CM-DiI (Molecular Probes C7000) was used to label choanocyte chambers in juveniles as previously described^16^, with slight modifications in the concentration used and incubation times. *A. queenslandica* juveniles were incubated in 1 μM CM-DiI in FSW for 30 minutes to 1 hour. This minimised the labelling of non-choanocyte cells. Despite this precaution, some non-choanocyte cells would be labelled in some individuals. Hence, all CM-DiI labelled juveniles were inspected by epifluorescence microscopy (Nikon Eclipse Ti microscope) immediately after CM-DiI was washed out, with juveniles detected with CM-DiI labelled cells outside of choanocyte chambers discarded from the study. Juveniles were allowed to develop for 0, 2, 4, 6, 12 or 24 hours post-incubation (hpi) with CM-DiI, then washed in FSW three times for 5 minutes and fixed^69^ without dehydration in ethanol. Fixed juveniles were washed three times in MOPST (1x MOPS buffer + 0.1% Tween). Nuclei were labelled with DAPI (1:1,000, Molecular Probes) for 30 minutes, washed in MOPST for 5 minutes and mounted using ProlongGold antifade reagent (Molecular Probes). All samples were observed using the ZEISS LSM 710 META confocal microscope, and image analysis was performed using the software ImageJ.

To visualise cell proliferation, the thymidine analogue EdU (Click-iT EdU AlexaFluor 488 cell proliferation kit, Molecular Probes C10337) was used as previously described^16,26^. To label S-phase nuclei, juveniles were incubated in FSW containing 200 μM EdU for 6 hours, washed in FSW and immediately fixed as described above. Fluorescent labelling of incorporated EdU was conducted according to the manufacturer’s recommendations prior to DAPI labelling and mounting in ProLong Gold antifade reagent as described above.

## Extended Data Figure Legends

**Extended Data Figure 1:**
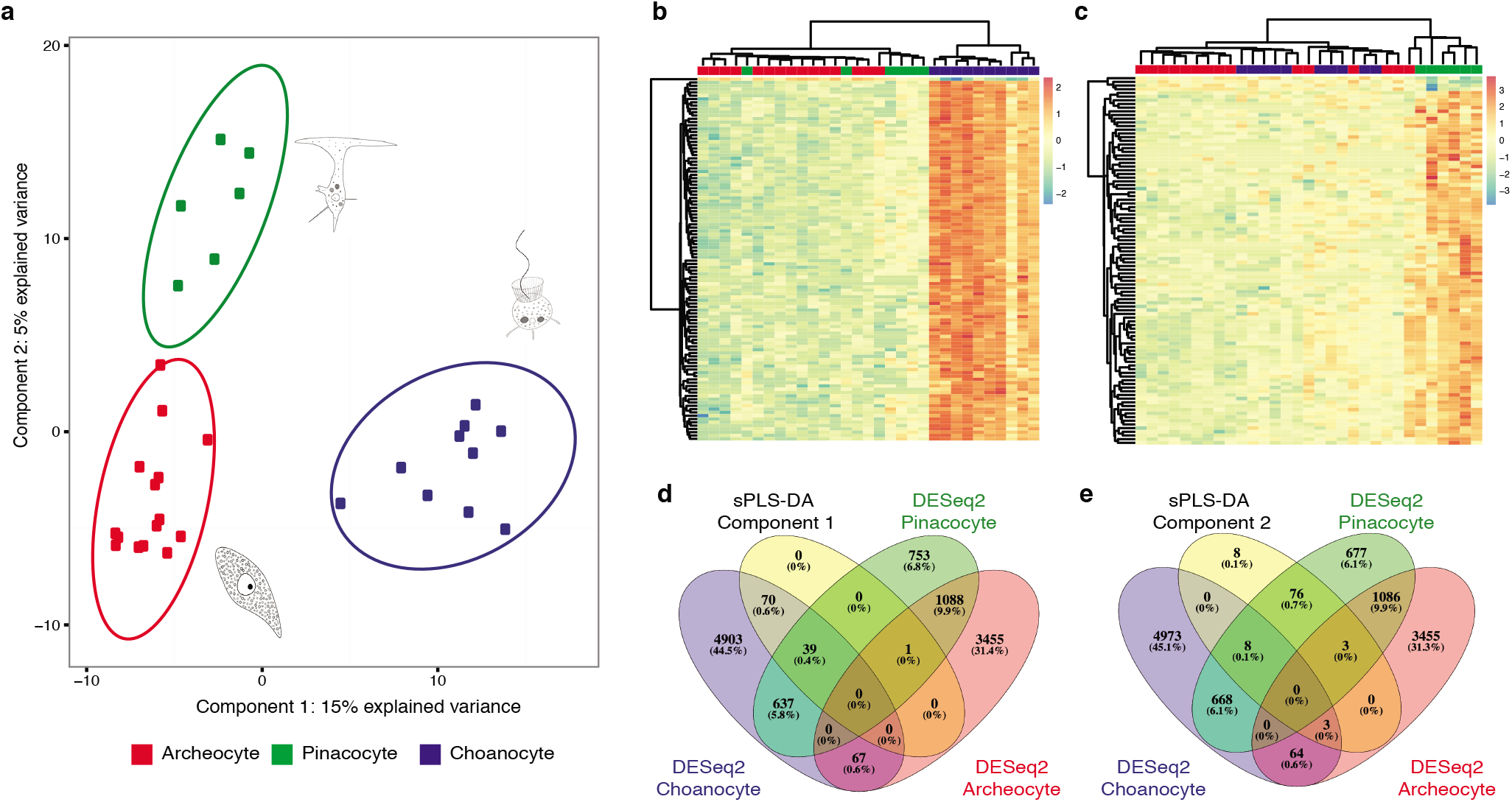
Sparse partial least squares discriminant analysis (sPLS-DA) of *Amphimedon queenslandica* choanocyte, archeocyte and pinacocyte transcriptomes. A supervised multivariate analysis, sPLS-DA, identified the gene models that best characterise differences in choanocytes (blue), archeocytes (red) and pinacocytes (green). **a,** Sample plot for the optimal number of gene models that discriminate cell types on the first two components; ellipses indicate 95% confidence intervals. **b, c,** Hierarchically-clustered heat maps show the expression of (b) the 110 gene models selected for the first component, and (c) the 98 gene models and 2 long non-coding RNAs selected for the second component, which accounted for 15% and 5% of explained variance, respectively. **d, e,** Venn diagrams summarise the significantly differentially expressed genes identified by the DESeq2 analyses, for each cell type, and the sPLS-DA on (d) the first and (e) the second sPLS-DA component. Percentages are of the total number of differentially expressed genes identified from all analyses.

**Extended Data Figure 2:**
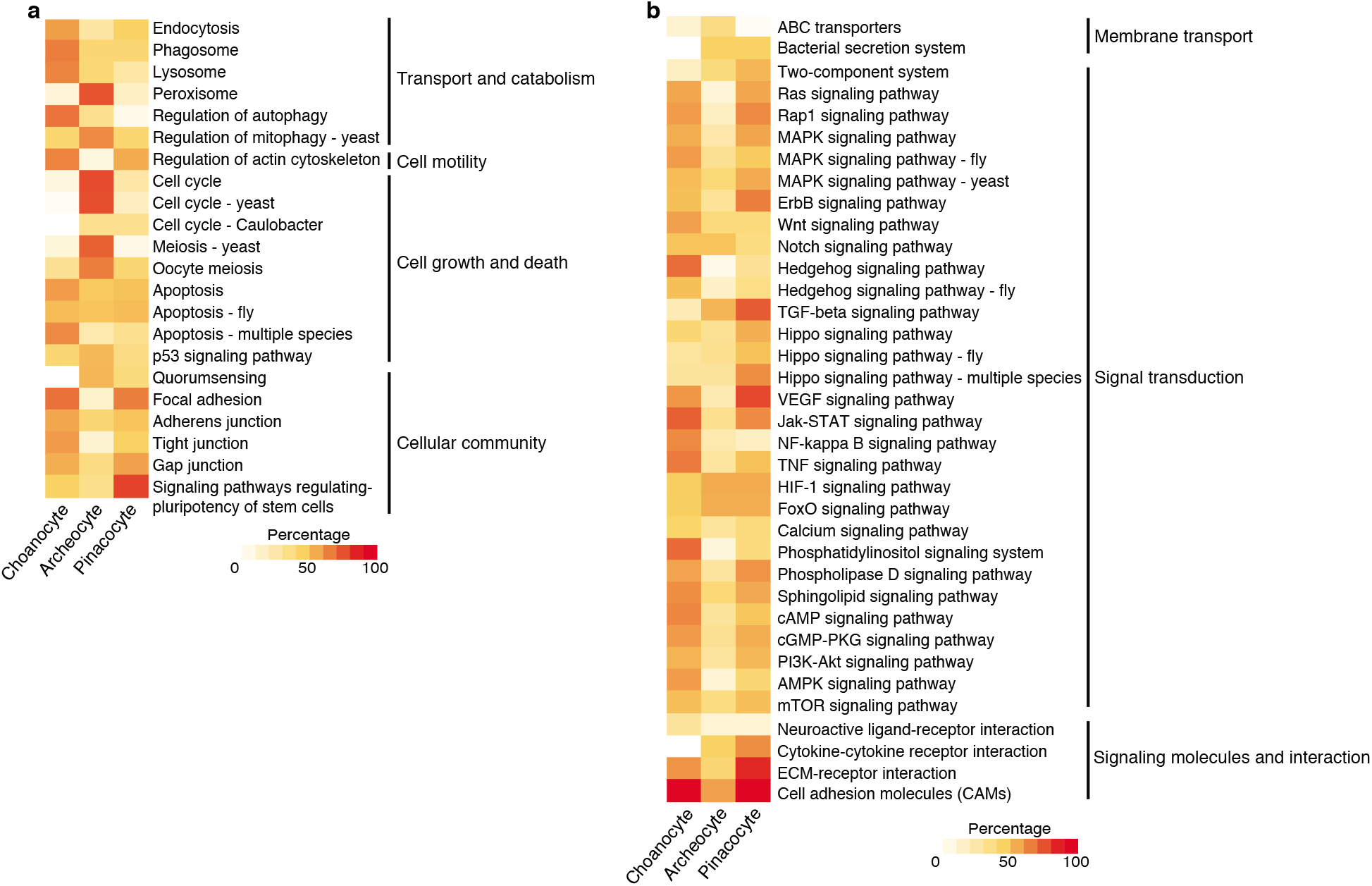
Percentage of KEGG cellular processes and environmental information processing (i.e. cell signalling) genes present in each cell type, corresponding to the number of components making up each KEGG category identified. **a,** Cellular processes genes. **b,** Environmental information processing (i.e. cell signalling) genes.

**Extended Data Figure 3:**
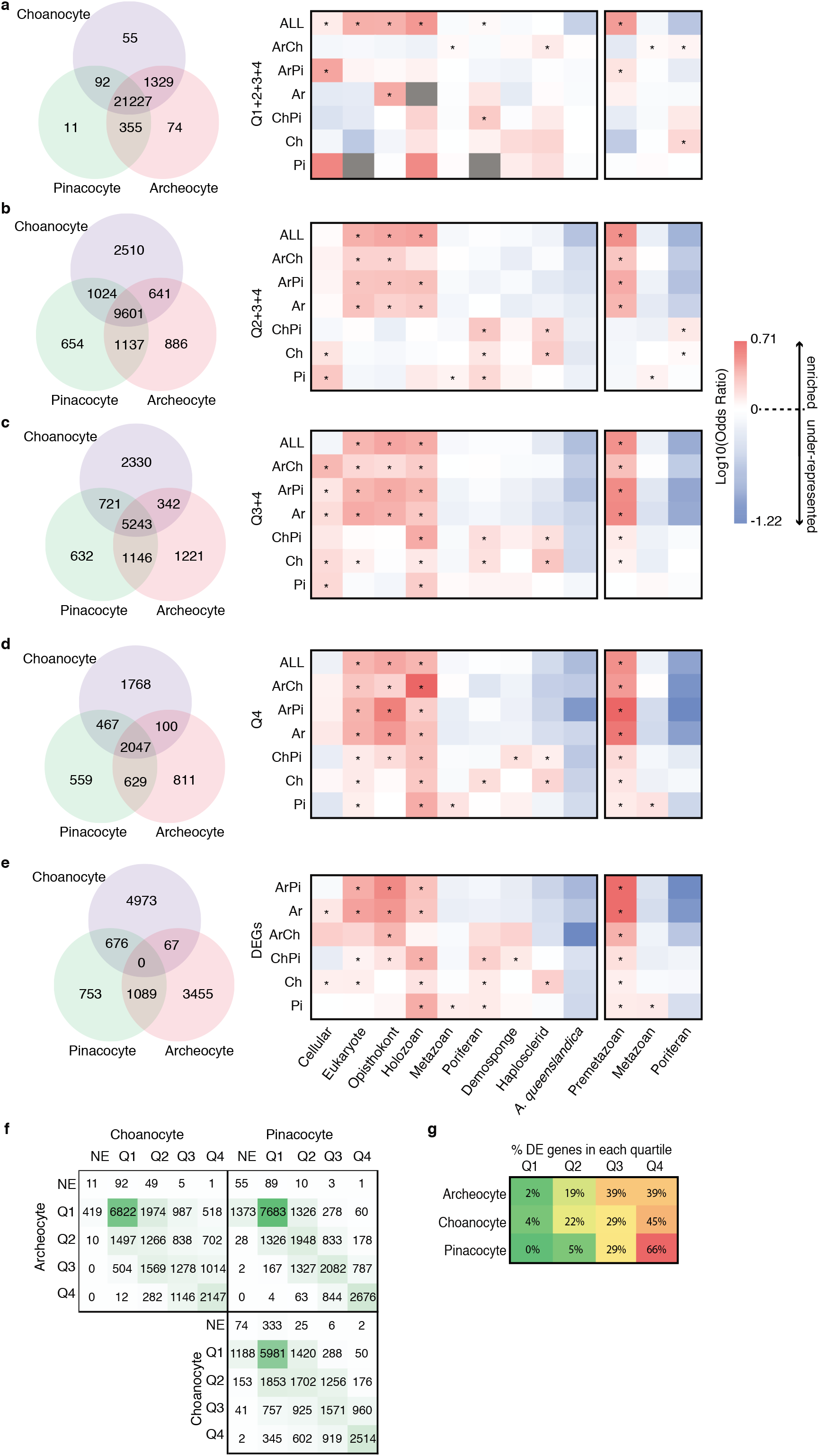
Evolutionary age of genes expressed in *Amphimedon queenslandica* choanocytes, archeocytes and pinacocytes using different expression thresholds. **a-e,** Phylostratigraphic enrichment of genes expressed in each cell type (Ar, archeocyte; Ch, choanocyte; Pi, pinacocyte; ArCh, archeocyte + choanocyte; ArPi, archeocyte + pinacocyte; ChPi, choanocyte + pinacocyte; ALL, all three cell types combined) at different expression thresholds. Expressed genes are parsed into quartiles based on transcript abundance in each of the cell types. Quartile 1 (Q1) includes the least abundant transcripts and Q4 the most abundant. a, Phylostratigraphy enrichment of all genes expressed in each of the cell types (i.e. Q1-Q4). **b,** Phylostratigraphy enrichment of genes expressed in the top three quartiles (i.e. excluding Q1). **c,** Phylostratigraphy enrichment of genes expressed in the top 50% (i.e. Q3 and Q4). **d,** Phylostratigraphy enrichment of the most highly expressed genes (i.e. Q4). **e,** For comparison, the evolutionary age of differentially expressed genes identified using differential expression analysis, DESeq2. Heat maps indicate enrichment (log odds ratio) of phylostrata contained in each gene list in comparison to the *A. queenslandica* genome. Asterisks mark significant (p < 0.05) enrichment. The heat maps on the far right are collapsed versions of the heat maps on the left, where the premetazoan category contains phylostrata from cellular to holozoan, and the poriferan category contains phylostrata from poriferan to *A. queenslandica*. To the left of each heat map is a Venn diagram, showing the number of genes in each cell type and sets of cell types. Grey boxes on the heat map indicate that there were no genes in that particular gene list characterised by the given phylostrata. **f,** Pairwise comparison illustrating the number of overlapping genes for each of the quartiles between the three cell types. The numbers in the cells are the number of genes common between two cell types (e.g. there are 1569 expressed genes in common between Q2 in choanocytes and Q3 in archeocytes). NE, not expressed. **g,** The percentage of differentially up-regulated genes identified in each of the cell types using DESeq2 in the four quartiles.

**Extended Data Figure 4:**
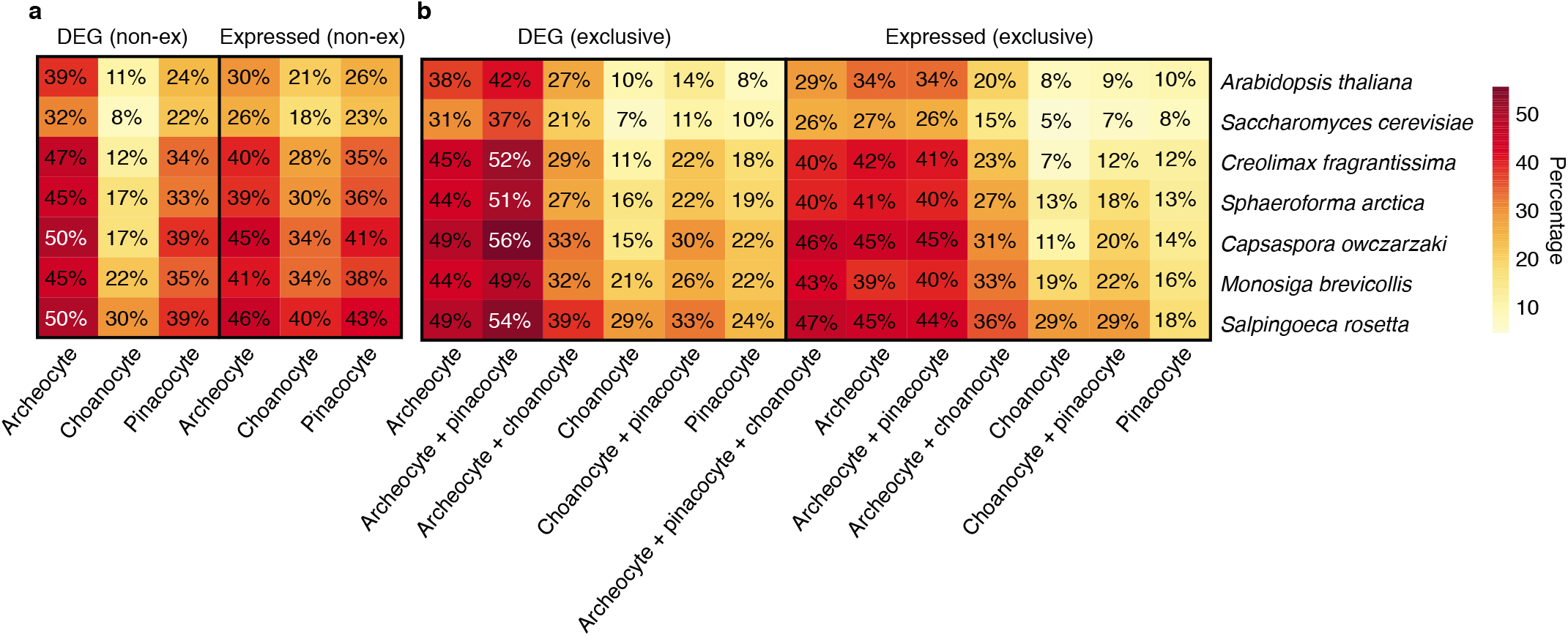
Orthologues shared between cell type-specific gene lists and non-metazoan eukaryotes. Heat map showing the percentage of *A. queenslandica* genes with orthogroups (OGs) shared with select eukaryotes. **a,** Percentage of genes with OGs shared between up-regulated and total expressed genes from non-exclusive lists (i.e. all genes expressed in each of the three cell types, not excluding genes that overlap between any two cell types). **b,** Percentage of genes with OGs shared between DEG and total expressed genes - exclusive lists (i.e. genes uniquely up-regulated or expressed in that cell type).

**Extended Data Figure 5:**
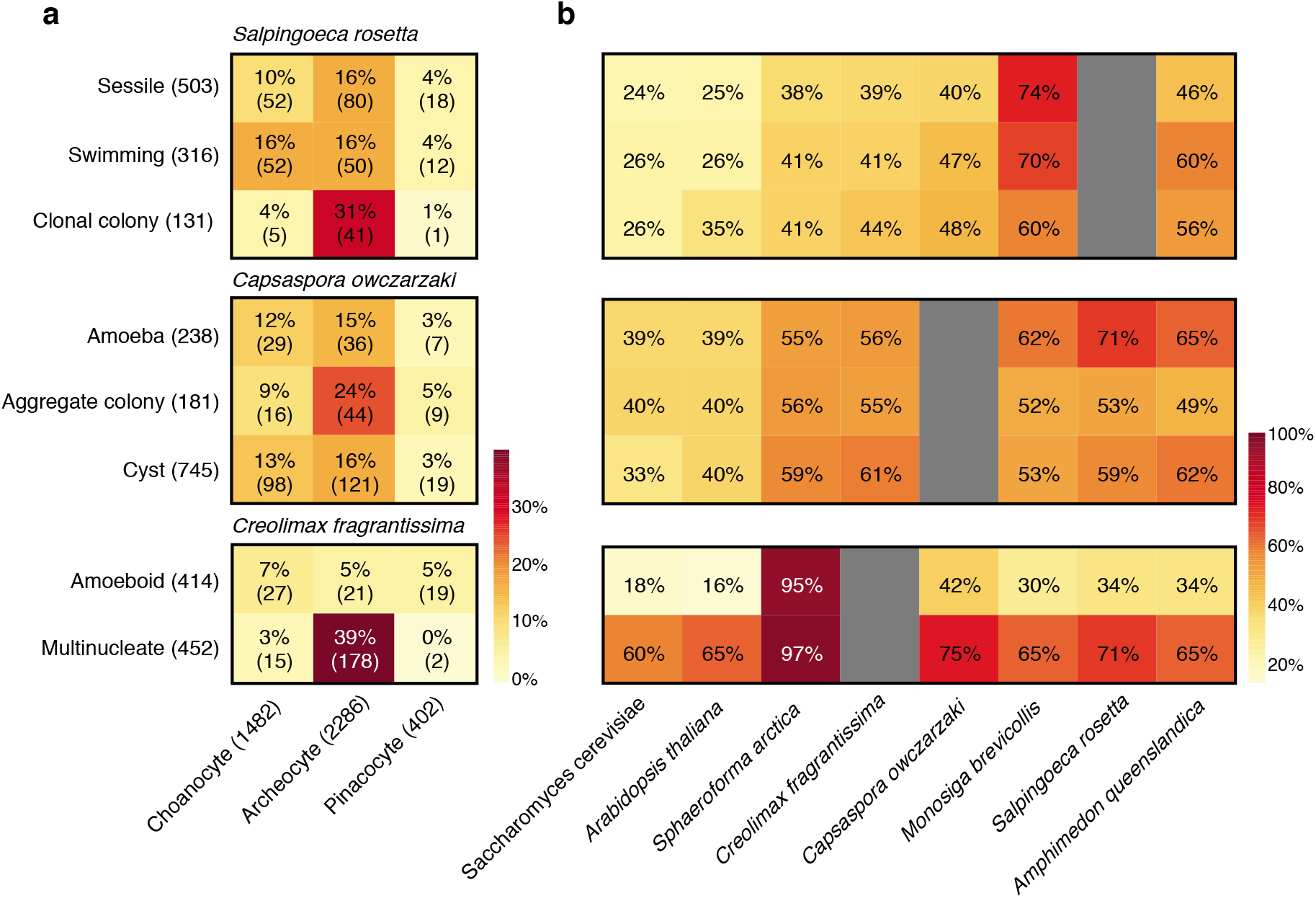
Orthologues found in *Salpingoeca rosetta, Capsaspora owczarzaki* and *Creolimax fragrantissima* life cycle stages, shared with *A. queenslandica* cell type transcriptomes and eukaryotic genomes. **a,** The percent and number (in parentheses) of differentially expressed OGs found in *Salpingoeca rosetta, Capsaspora owczarzaki* and *Creolimax fragrantissima* life cycle stages that are shared with *Amphimedon queenslandica* cell types. The numbers in parentheses alongside the unicellular holozoan cell states and sponge cell type names is the total number of OGs differentially expressed in that specific gene list. **b,** A heatmap showing the percentage of OGs shared between genes differentially expressed in *Salpingoeca rosetta, Capsaspora owczarzaki* and *Creolimax fragrantissima* life cycle stages, and genes present in other eukaryotic genomes.

**Extended Data Figure 6:**
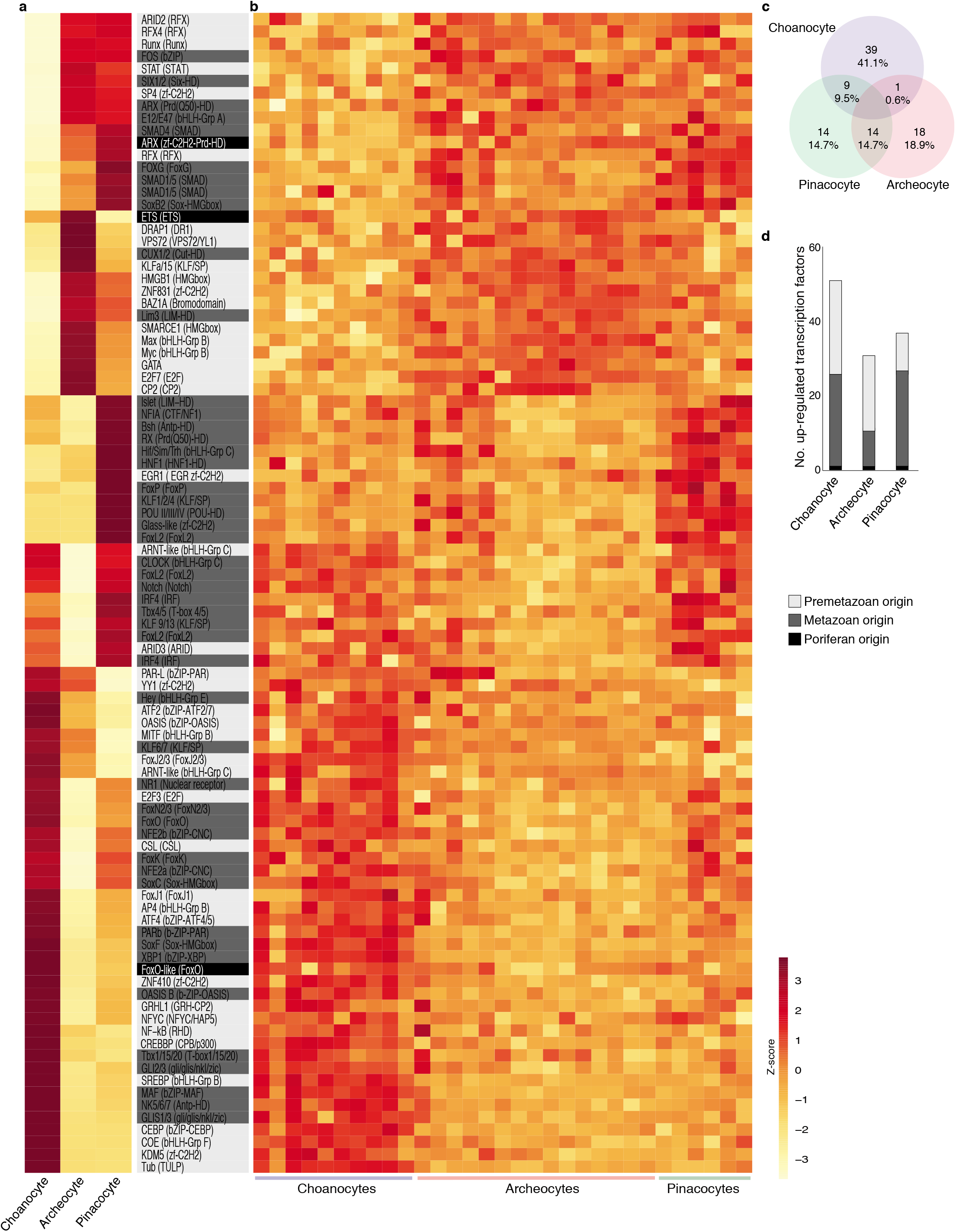
Heat map of transcription factor genes differentially expressed in choanocytes, archeocytes and pinacocytes. 94 transcription factor genes that are differentially expressed in *A. queenslandica* cell types are classified based on phylostratum: premetazoan (light grey); metazoan (dark grey; and poriferan (black). **a,** Heat map of expression levels in the three cell types combining all analysed CEL-Seq2 data. Gene names, families (in parentheses) and phylostrata shading are shown on the right. **b,** Heat map of expression levels of all CEL-Seq2 samples. Rows in b correspond to the rows and genes in a. **c,** Venn diagram summary of differentially up-regulated transcription factor genes between the three cell types using DESeq2. Percentages are of the total transcription factor genes differentially up-regulated in all cell types. **d,** Bar graph of the number and distribution of transcription factor genes based on evolutionary age in the three cell types.

**Extended Data Figure 7:**
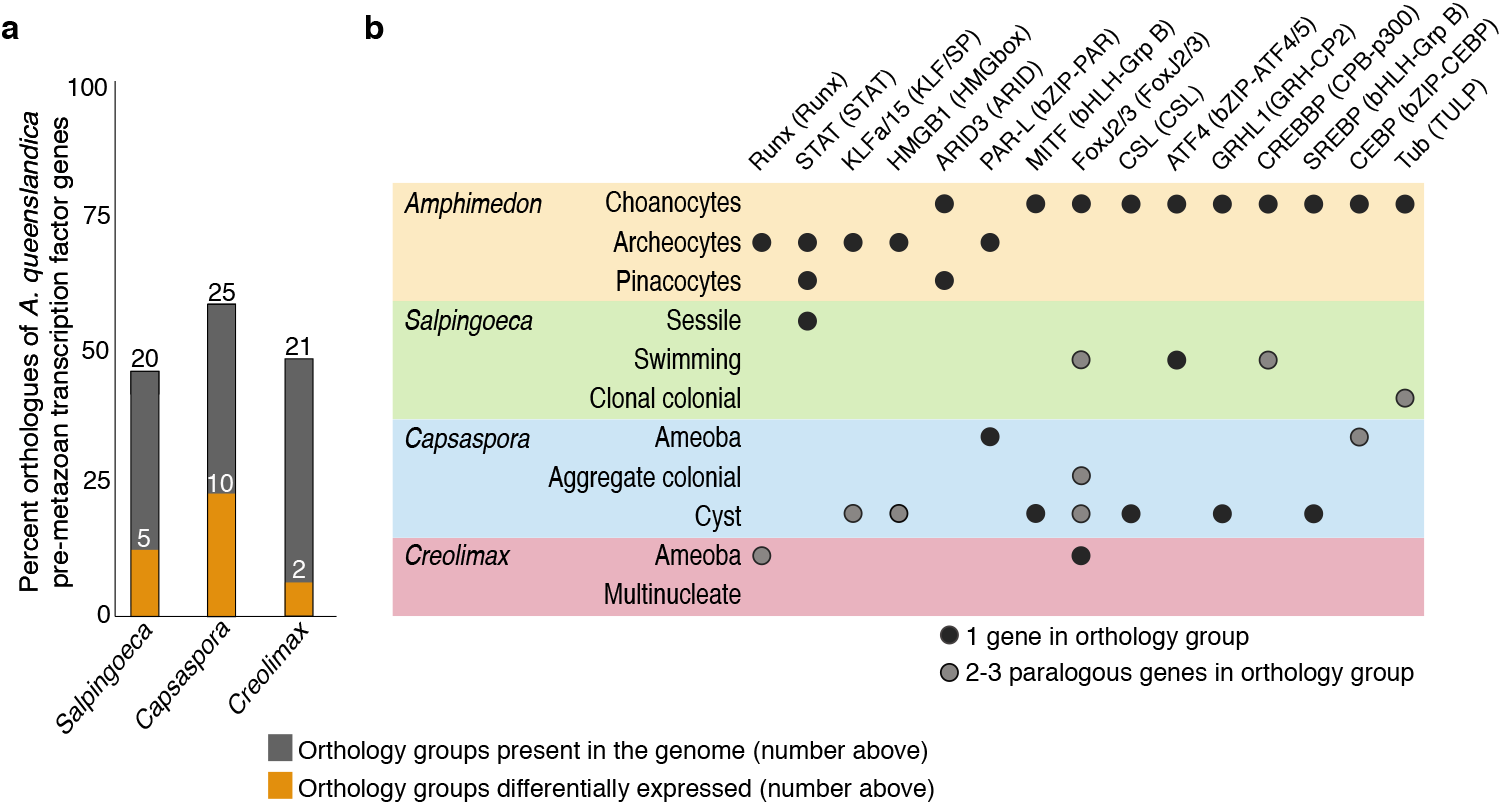
Analysis of premetazoan transcription factors in *Amphimedon* cells and unicellular holozoan cell states. **a,** The number and percentage of premetazoan transcription factor orthologues that are present in the genomes of *Salpingoeca rosetta, Capsaspora owczarzaki* and *Creolimax fragrantissima*. Percentages are based on the 43 premetazoan genes differentially expressed in the *A. queenslandica* cell types (Extended Data Fig. 5). The number of transcription factor orthologues in the genome is listed above the bar. The orange bar depicts the percent and number of unicellular holozoan premetazoan transcription factor orthologues that are significantly differentially up-regulated in at least one cell state. **b,** The 15 premetazoan transcription factor orthology groups (listed along the top) that are significantly up-regulated in at least one *Amphimedon* cell type and one unicellular holozoan cell state. Dots correspond to the cell types and states this occurs. Black dots, orthology group with one gene member; grey dots, orthology group comprised of two of more paralogues (see Supplementary File S8 for details).

**Extended Data Figure 8:**
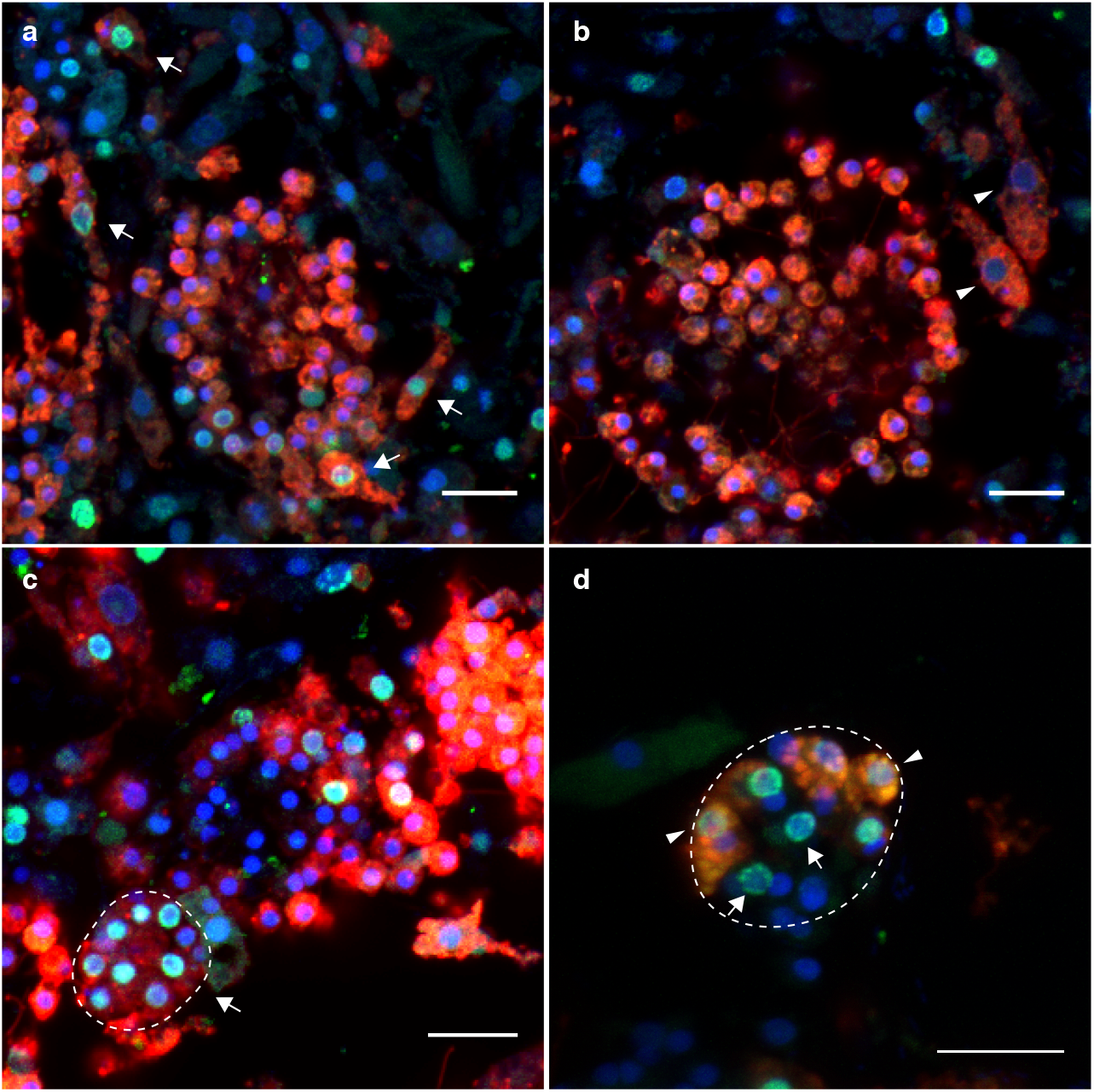
Choanocyte dedifferentiation into an archeocyte does not require cell division. **a, b,** 4 day old juveniles 6 hours after CM-DiI and EdU labelling. **a,** CM-DiI labelled archeocytes with EdU incorporation (arrows) found near choanocyte chambers. **b,** Labelled archeocytes without EdU incorporation (arrowheads), indicating dedifferentiation from choanocytes without cell division. Scale bars: 10 μm. **c, d,** Choanocyte-derived archeocytes are capable of generating new choanocyte chambers. **c,** 4 day old juvenile 6 hours after CM-DiI and EdU labelling. Early choanocyte chamber (dotted line) completely labelled with CM-DiI and EdU, indicating CM-DiI labelled archeocytes, with large nuclei, are forming this chamber. The absence of cilia and space at the center of this structure indicates it is not yet a functional choanocyte chamber. **d,** 4 day old juvenile 12 hours after CM-DiI and 6 hours after EdU labelling. Early choanocyte chamber (dotted line) with multiple EdU labelled cells, with both CM-DiI labelled choanocytes (arrowheads) and non-CM-DiI labelled choanocytes (arrows) indicate multiple cell lineages contributing to the formation of this chamber. Scale bars: **a-d,** 10 μm.

## Supplementary Video Legends

**Supplementary Video 1: Time-lapse video of a 4 day old juvenile *Amphimedon queeslandica*.** This 10 second video captures 20 minutes of cell behavior on the outer edge of the juvenile. Annotated are (i) a choanocyte chamber (cc) comprising of multiple tethered choanocytes, (ii) three migrating archeocytes (ar) – there are multiple other archeocytes in this video, and (iii) a pinacocyte (pi), which comes in and out of focus and is characterised by a thin, transparent cytoplasm with small refractive vesicles. Scale bar, 10 μm.

**Supplementary Video 2: Capture and washing of a dissociated archeocyte.** All cells and choanocyte chambers used in this study were fixed or frozen in less than 15 minutes after dissociation from the intact sponge.

**Supplementary Video 3: Time-lapse video of choanocytes transdifferentiating and evacuating chambers in 4 day old juvenile *Amphimedon queeslandica*.** Matching 8-second videos captures 120 min of CM-DiI labelled choanocytes (left, red; right, white), which are initially located in distinct chambers (arrows on four chambers), undergoing transdifferentiation and migrating from the chambers. At the end of the video, none of the CM-DiI labelled chambers are recognisable. Note that many cells vacating the choanocyte chambers are larger, consistent with choanocytes dedifferentiating into larger archeocytes. Scale bar, 40 μm.

## Supplementary Files

**Supplementary File S1. The counting report of the cell type specific CEL-Seq2 samples**

**Supplementary File S2. Table summarising the details and the statistics of the demultiplexing and mapping steps for the cell type specific CEL-Seq2 samplesE**⍰

**Supplementary File S3. Table of differentially expressed gene lists from DESeq2 with BLAST2GO annotations and phylostrata ID**⍰

**Supplementary File S4. Table of differentially expressed gene lists from sPLS-DA with BLAST2GO and KEGG annotations and phylostrata ID**⍰

**Supplementary File S5. Table of KEGG enrichment analysis results on differentially expressed gene lists from DESeq2** This spreadsheet contains the output of KEGG enrichment analyses performed on each cell type DEG list. The first sheet contains percentage values of genes/components identified in the DEG lists relative to the *A. queenslandica* genome.

**Supplementary File S6. Table of the phylostrata enrichment of the differentially expressed gene lists from DESeq2**⍰

**Supplementary File S7. Table of cell-type gene lists and transcription factor lists from the quartile expression analyses**

**Supplementary File S8. Table of transcription factor genes expressed in the three cell types and in the differentially expressed gene lists from DESeq2**⍰

**Extended Data Table 1.**
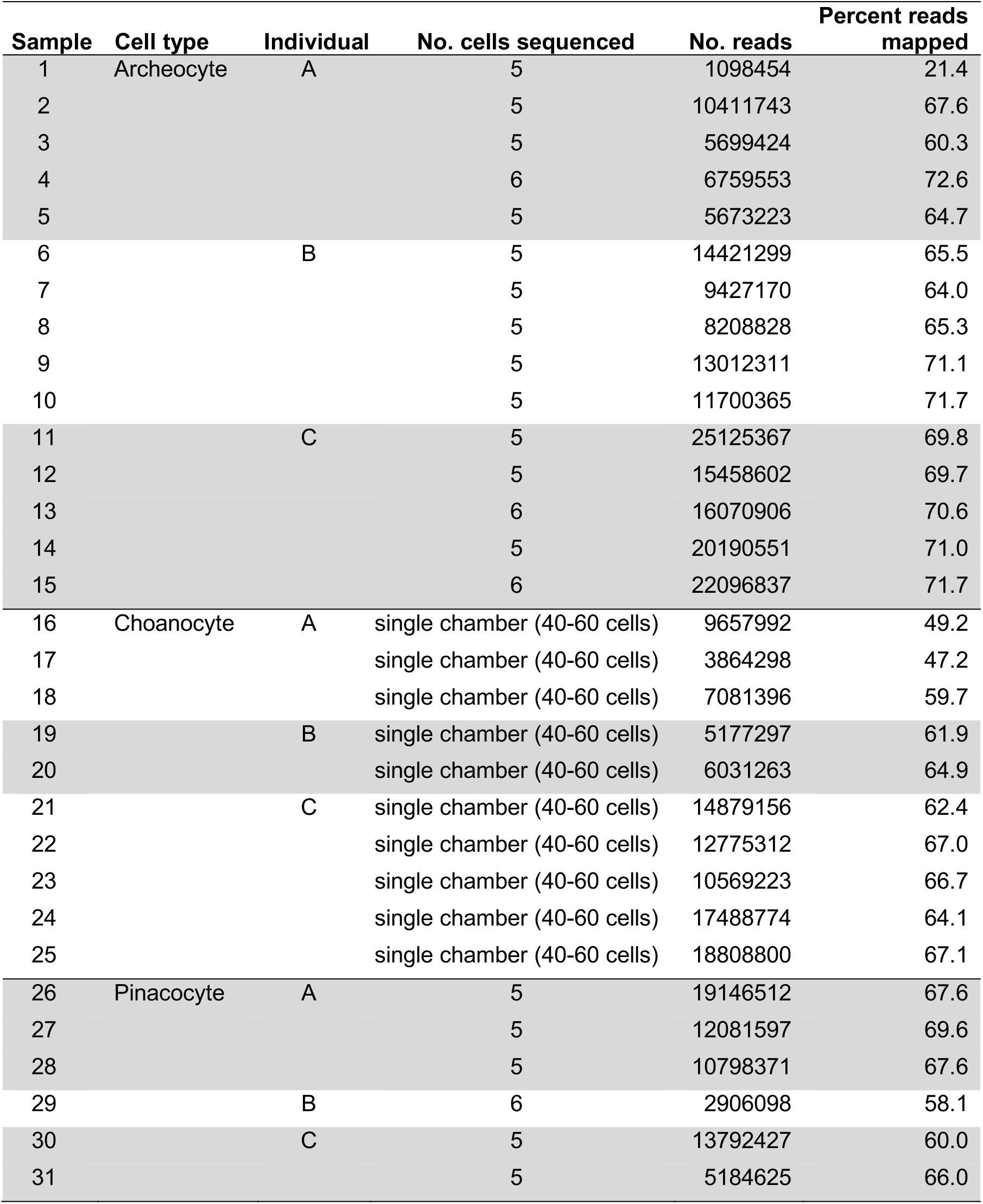
Summary of CEL-Seq2 samples used in this study

| Sample | Cell type | Individual | No. cells sequenced | No. reads | Percent reads mapped |
| --- | --- | --- | --- | --- | --- |
| 1 | Archeocyte | A | 5 | 1098454 | 21.4 |
| 2 |  |  | 5 | 10411743 | 67.6 |
| 3 |  |  | 5 | 5699424 | 60.3 |
| 4 |  |  | 6 | 6759553 | 72.6 |
| 5 |  |  | 5 | 5673223 | 64.7 |
| 6 |  | B | 5 | 14421299 | 65.5 |
| 7 |  |  | 5 | 9427170 | 64.0 |
| 8 |  |  | 5 | 8208828 | 65.3 |
| 9 |  |  | 5 | 13012311 | 71.1 |
| 10 |  |  | 5 | 11700365 | 71.7 |
| 11 |  | C | 5 | 25125367 | 69.8 |
| 12 |  |  | 5 | 15458602 | 69.7 |
| 13 |  |  | 6 | 16070906 | 70.6 |
| 14 |  |  | 5 | 20190551 | 71.0 |
| 15 |  |  | 6 | 22096837 | 71.7 |
| 16 | Choanocyte | A | single chamber (40-60 cells) | 9657992 | 49.2 |
| 17 |  |  | single chamber (40-60 cells) | 3864298 | 47.2 |
| 18 |  |  | single chamber (40-60 cells) | 7081396 | 59.7 |
| 19 |  | B | single chamber (40-60 cells) | 5177297 | 61.9 |
| 20 |  |  | single chamber (40-60 cells) | 6031263 | 64.9 |
| 21 |  | C | single chamber (40-60 cells) | 14879156 | 62.4 |
| 22 |  |  | single chamber (40-60 cells) | 12775312 | 67.0 |
| 23 |  |  | single chamber (40-60 cells) | 10569223 | 66.7 |
| 24 |  |  | single chamber (40-60 cells) | 17488774 | 64.1 |
| 25 |  |  | single chamber (40-60 cells) | 18808800 | 67.1 |
| 26 | Pinacocyte | A | 5 | 19146512 | 67.6 |
| 27 |  |  | 5 | 12081597 | 69.6 |
| 28 |  |  | 5 | 10798371 | 67.6 |
| 29 |  | B | 6 | 2906098 | 58.1 |
| 30 |  | C | 5 | 13792427 | 60.0 |
| 31 |  |  | 5 | 5184625 | 66.0 |

